# Rapid optimization of protein function in mammalian cells via microbe-independent deep assembly and screening

**DOI:** 10.1101/2025.03.30.646252

**Authors:** Yan Wu, Pengli Wang, Lan Xiang Liu, Chao Gao, Qin Qin, Matt Hageman, Thomas A. Kirkland, Yichi Su, Michael Z. Lin

## Abstract

Random mutagenesis and deep mutational scanning (DMS) are widely used to optimize proteins by oversampling large libraries in microbial cells, selecting cells expressing favorable variants, and sequencing to identify enriched variants. However, these methods are slow and costly, require effort to establish selection methods for a given protein function, and do not yield data on lower-performing variants. Here, we describe Microbe-Independent Deep Assembly and Screening (MIDAS), a rapid, high-throughput method for optimizing protein function directly in mammalian cells. As a demonstration, we applied MIDAS to improve a newly designed neurotransmitter bioluminescent indicator (NeuBI) for acetylcholine (ACh). MIDAS systematically optimized interdomain linkers, identified mutational hotspots, and exhaustively scanned amino acid combinations, in each case relating specific sequences to protein performance. MIDAS-optimized variants exhibited improved performance in vivo, highlighting the potential of MIDAS for improving protein function in mammalian systems.

## Introduction

Methods to accelerate the functional optimization of proteins in mammalian cells would be highly desirable, especially for the development of synthetic protein switches such as biosensors and ligand-responsive controllers, which often are suboptimal in output activity or in responsiveness to input. While machine learning algorithms have made large strides in solving the problem of designing primary sequences to create desired three-dimensional folds or protein-protein interfaces, these algorithms lack training for optimizing the transduction of conformational changes from input regions to modulate output functions. In addition, even for the well-defined static problem of optimizing ligand affinity, current machine learning algorithms do not reliably identify the most optimal protein sequence or small sets of near-optimal sequences. Rather they typically generate hundreds of candidates which then need to be generated and screened, typically via multi-well or brute-force molecular cloning methods.

The process of optimizing dynamic proteins for function in mammalian cells generally involves performing steps in both microbial and mammalian cells. Mammalian validation is critical, as some mutations that improve performance in bacteria may prove ineffective in mammalian systems^1^, and binding affinities of proteins to small molecules can differ substantially between bacterial and mammalian settings^2^. Libraries can be expressed and screened in bacterial or yeast cells for higher throughput and shorter time to initial selection. When cells are enriched into pools rather than into clones, and high-throughput sequencing is applied to these pools, this method is referred to as deep mutational scanning (DMS)^3^. Regardless, with microbial-based screens, validation of protein performance in mammalian cells becomes a time-consuming step, involving isolation of expression plasmids, plasmid preparation, transfection, and reassessment of protein function^2,4,5^.

Multiple methods have been established for screening for improved protein function in mammalian cells^6^, but they still involve preceding library preparation steps in bacteria. For functions that can be assessed in mammalian cells with an optical readout, screening can be performed by imaging in mixed mammalian cultures and retrieved by mechanical picking^7^, or by fluorescence-activated cell sorting (FACS) which can be combined with next-generation sequencing in a DMS workflow^8^, but at the cost of noise from cell-cell variability. Practically, multiple rounds of enrichment and/or large secondary screens^9–11^, or repeated experiments in the case of DMS^8^, are often required to identify improvements with high confidence. For the most reliable functional selections, mammalian cells expressing mutant libraries can be grown from single cells seeded in multiwell plates and assayed as populations, achieving medium throughput (10^4^ per day). To introduce variants into mammalian cells, these strategies utilize lentiviral transduction at limiting doses or recombinase-mediated insertion of transgenes into single genomic sites^12^. However, obtaining the library DNA for either lentiviral packaging or in-cell recombination still requires prior plasmid construction and purification in bacteria. One method that does not require bacterial library generation involves selective propagation of viruses encoding improved proteins in a continually mutagenic environment, but this strategy is limited to protein functions that can be linked to the transcription of viral replication genes^13,14^. Thus, existing generalizable strategies for optimizing protein function in mammalian cells involve weeks of work on library generation in microbial hosts and characterization in mammalian cells.

If a method existed for performing library construction and screening entirely in mammalian cells without a microbial step, then optimizing protein functions for mammalian performance could be greatly accelerated. As a test case for a new technique for protein optimization, we chose to develop a bioluminescent indicator for the neurotransmitter acetylcholine (ACh). The study of ACh function in normal physiology and its dysregulation in disease would benefit from a technology for visualizing ACh using bioluminescence. ACh plays multiple essential roles in the nervous systems of animals. In vertebrates, it serves as the primary neurotransmitter at neuromuscular junctions and in the parasympathetic and enteric nervous systems^15,16^. In the mammalian central nervous system (CNS), ACh functions as a neuromodulator involved in arousal, attention, learning, and memory^17,18^. Cholinergic neurons, originating from the nucleus basalis, project throughout the cortex, with the highest innervation densities in the hippocampus. Reduced cholinergic activity occurs early in several neurodegenerative disorders, including Alzheimer’s disease^19,20^, Lewy body dementia^21^, and Parkinson’s disease^22^. In Parkinson’s disease, dysautonomia due to degeneration of cholinergic parasympathetic neurons in the vagus nerve is also commonly observed^23^.

While fluorescent indicators for ACh have been developed^5,24^, their applications in vivo necessitate the implantation of optical elements such as fibers, windows, or lenses. While feasible in the brain due to the essentially non-motile nature of the brain and surrounding skull^24–26^, implantation of optical elements is less compatible with visualizing tissues or organs in other parts of the body. In small animal models, a bioluminescent indicator would allow noninvasive, real-time monitoring of ACh release in locations where optical elements cannot be fixed while using affordable, readily available equipment. However, no bioluminescent indicators for ACh have been developed to date. In the absence of a pre-existing bioluminescent neurotransmitter indicator design that could be adapted, we anticipated that developing an ACh indicator would require extensive optimization to achieve sufficient responsiveness for detecting physiological changes in ACh levels in vivo.

Here, we report the development of microbe-independent deep assembly and screening (MIDAS) as a method for protein optimization in mammalian cells. MIDAS integrates three steps in rapid succession: multi-well PCR to construct libraries of all possible variants at targeted sites, direct transfection of linear PCR products into mammalian cells, and cell-based phenotypic screening. In addition, it can deeply sample machine learning-identified mutational hotspots by producing and screening each allelic variant exactly once, maximizing efficiency. Using MIDAS, we enhanced the output dynamic range (overall responsivity) of a prototype Acetylcholine Neurotransmitter Bioluminescence Indicator (ACh-NeuBI) by 29-fold in just five rounds of mutagenesis. The optimized ACh-NeuBI successfully detected exogenous ACh noninvasively in vivo, with up to 100-fold responses.

## Results

### Design of a prototype acetylcholine bioluminescent indicator

We chose to create a bioluminescent small-molecule indicator based on a periplasmic binding protein (PBP) as a representative test case for comprehensive protein optimization. Bacterial PBPs are bi-lobed proteins that bind to a variety of small molecules^27,28^, and have been used as a template for fluorescent indicators of glutamate, acetylcholine, serotonin, and GABA^2,5,26,29^. In these fluorescent indicators, a circularly permuted GFP is inserted between the N-terminal (NT) and C-terminal (CT) lobes. Conformational changes in the PBP upon ligand binding are transduced to GFP side chain conformational changes to induce chromophore deprotonation^2^. So far, however, there is no bioluminescent indicator for small molecule analytes based on PBPs. Given the versatility of PBPs in binding a large range of analytes of importance in mammalian physiology and metabolism, and the general utility of bioluminescence for non-invasive imaging in the body^30–32^, we set out to create a PBP-based bioluminescent indicator.

We selected ACh for detection by a PBP-based bioluminescent indicator, as cholinergic signaling regulates motor and metabolic functions throughout the mammalian body and is disrupted in various medical disorders, including autoimmune and degenerative conditions. We hypothesized that we could design a bioluminescent indicator of ACh using NanoLuc and a mutant OpuBC domain from *Thermoanaerobacter sp. X513* previously engineered for ACh specificity over choline^5^ (**Fig. 1a**). To first identify the optimal topology of fusions of NanoLuc and OpuBC fragments, we conducted a screening of various forms of circularly permuted NanoLuc (cpNanoLuc), including 64/65, 131/132, 154/155, and LgBiT/SmBiT^33^ (**Fig. S1a)**. Each construct was cloned and expressed in HEK293A cells using transient transfection, and their luminescence was measured before and after treatment with 100 μM ACh. Only the insertion between LgBiT and SmBiT showed an increase in luminescence upon ACh treatment, of 22% (**Fig. S1b**). This result was not surprising, as assembly of LgBiT and SmBiT can be robustly modulated by intramolecular conformational changes^31,32^. We designated this construct as Acetylcholine Neurotransmitter Bioluminescent Indicator 0.1 (ACh-NeuBI0.1) and selected it for further optimization.

**Fig. 1.**
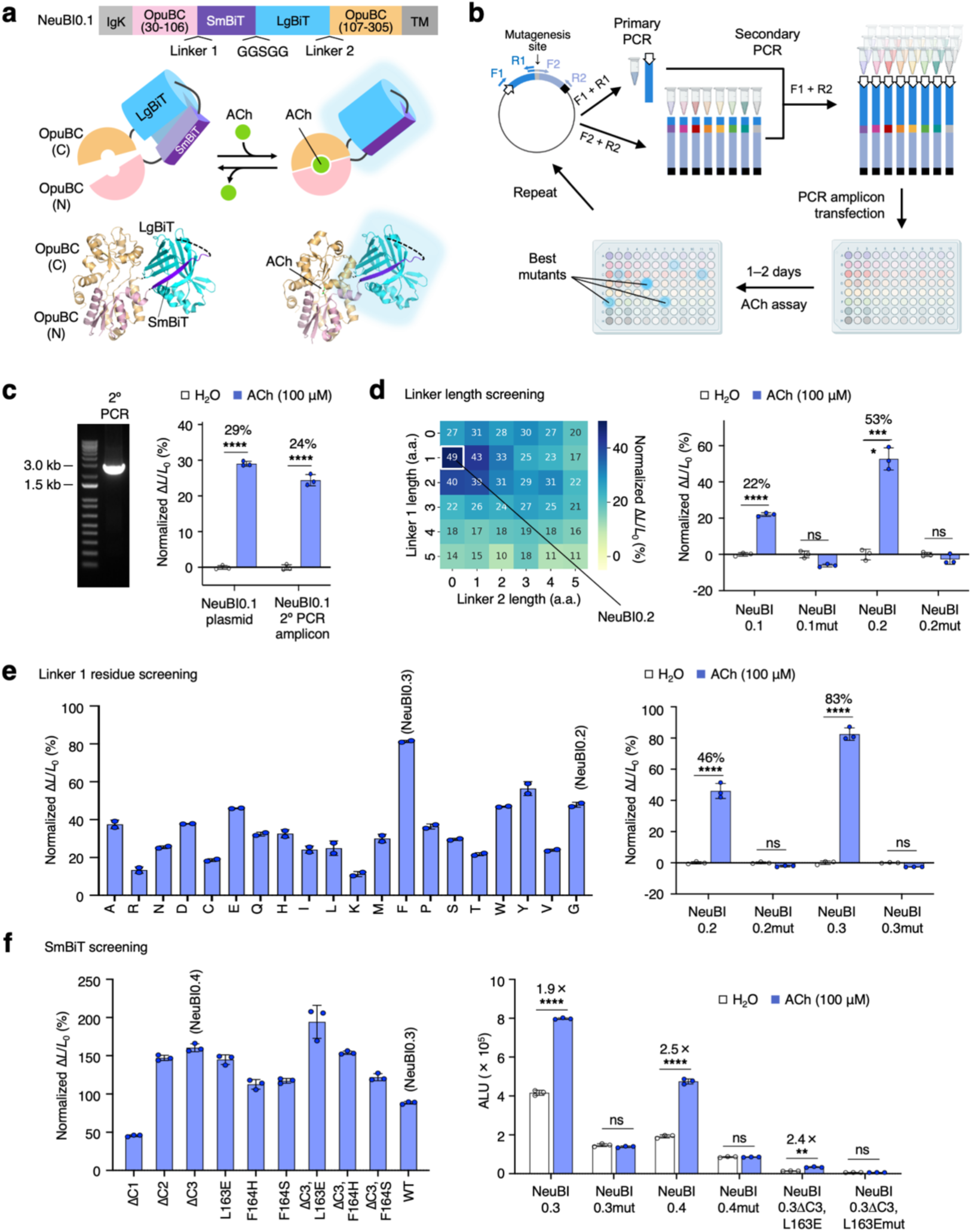
Design and initial optimization of ACh-NeuBI using MIDAS. **a**, Domain arrangement, cartoon, and structural model of ACh-NeuBI0.1 rendered from PDB coordinates 6URU (OpuBC) and 5IBO (NanoLuc). Acetylcholine (ACh) binding to OpuBC induces complementation of circularly permutated NanoLuc (cpNanoLuc) and thereby increased bioluminescence. **b**, MIDAS strategy. Primary polymerase chain reactions (PCRs) amplify overlapping fragments of a full-length gene including promoter (white arrow) and polyadenylation signal (black box). The overlap reason is selected to include the site targeted for mutagenesis so that one of the overlapping primers can be purchased in multi-well format to create all desired mutation deterministically with the aid of multi-well PCR machines. The secondary PCR amplicons are transfected into HEK cells for testing 24–48 h later. **c**, Validation of PCR amplicon transfection. Left, gel electrophoresis of ACh-NeuBI0.1. PCR yields a clean 2.6-kb band as expected. Right, responses of ACh-NeuBI0.1 expressed in HEK293A cells by plasmid transfection or that expressed by PCR products transfection to ACh (100 μM). **d**, Linker length screening by MIDAS. Left, heatmap showing the responses of the screened variants to ACh (100 μM). Right, responses of plasmid-transfected ACh-NeuBI0.1, ACh-NeuBI0.1mut, ACh-NeuBI0.2, and ACh-NeuBI0.2mut to ACh (100 μM) in HEK293A cells. **e**, Linker 1 residue screening. Left, responses of PCR product-transfected ACh-NeuBI0.2 mutants with different amino acids on linker 1 to ACh (100 μM). Right, responses of plasmid-transfected ACh-NeuBI0.2, ACh-NeuBI0.2mut, ACh-NeuBI0.3, and ACh-NeuBI0.3mut to ACh (100 μM) in HEK293A cells. **f**, SmBiT screening. Left, responses of PCR product-transfected ACh-NeuBI0.3 mutants with different SmBiT sequences to ACh (100 μM). Right, responses of plasmid-transfected ACh-NeuBI0.3, ACh-NeuBI0.3mut, ACh-NeuBI0.4, ACh-NeuBI0.4mut, ΔC3, L163E, and ΔC3, L163E mut to ACh (100 μM) in HEK293A cells. ALU, arbitrary luminescence units. **c–f**, ns, not significant; *, p < 0.05; **, p < 0.01; ***, p < 0.001; **** p < 0.0001, by unpaired two-tailed Student’s t-test compared to the H_2_O control. Error bars, SD.

### Linker optimization in mammalian cells by MIDAS

We first sought to optimize the linker lengths between OpuBC (30-106) and SmBiT (linker 1) and between LgBiT and OpuBC (107-305) (linker 2). To bypass the labor-intensive and time-consuming steps of plasmid cloning in bacteria, we hypothesized that we could accelerate the expression and screening of sensor variants in mammalian cells by constructing entire genes with the desired modifications as linear PCR products in multi-well blocks and directly transfecting them into mammalian cells in matching multi-well plates. Variants could then be directly screened for improved function in their desired cell type and subcellular location (**Fig. 1b**). We refer to this method as Microbial-Independent Deep Assembly and Screening (MIDAS). MIDAS is a faster and cost-effective alternative to traditional cloning methods, as it bypasses the time-consuming and expensive cloning steps of ligation or recombineering, bacterial transformation, bacterial growth, plasmid purification, and sequence verification. To validate the strategy, we transfected HEK293A cells with PCR-amplified genes encoding surface-anchored ACh-NeuBI0.1 fused to the PDGFR transmembrane domain, including a promoter and polyadenylation signal, and compared responses to ACh stimulation compared with transfection of ACh-NeuBI0.1-expressing plasmid. We found that responses were similar between PCR amplicon-transfected and plasmid-transfected cells (**Fig. 1c**).

To optimize linker lengths between cpNanoLuc and the OpuBC fragments, we designed and screened linkers consisting of zero to five glycine residues on each end. Using MIDAS, we constructed and screened all 25 possible combinations in HEK293A cells without cloning. Within these combinations, whose responses exhibited more than a 4-fold difference from weakest to strongest, we identified 1 and 0 amino acids at linkers 1 and 2 to be optimal, supporting a ∼50% increase in luminescence following treatment with 100 μM ACh (**Fig. 1d,** left). We designated this optimized variant as ACh-NeuBI0.2, and confirmed its performance after traditional plasmid cloning and transfection (**Fig. 1d**, right). For control experiments, we introduced a mutation (Y140A)^5^ at the ACh-binding site of OpuBC into ACh-NeuBI0.1 and ACh-NeuBI0.2. As expected, the non-binding ACh-NeuBI mutants (ACh-NeuBI0.1mut and ACh-NeuBI0.2mut) showed no signal increase upon ACh treatment (**Fig. 1d**, right), confirming that the responses observed in ACh-NeuBI0.1 and ACh-NeuBI0.2 were specifically triggered by ACh binding.

Next, to identify the optimal residue for linker 1 in ACh-NeuBI0.2, we systematically tested all 20 amino acids using MIDAS. Phenylalanine (F) emerged as the most effective, improving the response to ∼80% (**Fig. 1e**, left). This variant was designated as ACh-NeuBI0.3 and validated by traditional cloning and plasmid transfection (**Fig. 1e**, right). Subsequently, within ACh-NeuBI0.3, we employed MIDAS to screen SmBiT variants with varied affinities to LgBiT^31^. Notably, both SmBiT (ΔC3) and SmBiT (ΔC3, L163E) exhibited substantial enhancement in their responsiveness (**Fig. 1f**, left). Upon revalidation, both showed a ∼2.5-fold increase upon 100 μM ACh exposure, while their non-binding mutants showed no significant signal change (**Fig. 1f**, right). However, the version with SmBiT (ΔC3, L163E) was extremely dim. Due to its superior brightness, the variant with SmBiT (ΔC3) was designated ACh-NeuBI0.4 and selected for further development.

### Red-shifting ACh-NeuBI emission via RET

The emission peak of NanoLuc at ∼450 nm is not ideal for tissue penetration due to significant absorption in biological tissues of light at wavelengths shorter than 600 nm^34^. Thus, to improve the retrieval of emitted light through tissue, we investigated attaching a fluorescent protein to ACh-NeuBI.0.4 to shift a fraction of emission to wavelengths longer than 600 nm via resonance energy transfer (RET) (**Fig. S2a**). Specifically, we tested CyOFP1, which features an excitation spectrum that overlaps favorably with NanoLuc’s emission spectrum, as well as the standard fluorescent proteins mScarlet-I and mScarlet3 (**Fig. S2b**). Both fluorescent proteins emit with a peak wavelength below 600 nm and with the majority of emitted photons below 620 nm, and are thus most appropriately considered orange fluorescent proteins^35^. Inserting a tandem dimer of CyOFP1 between SmBiT and LgBiT (B1), an arrangement topologically resembling the high-RET Antares luciferase, produced no RET (**Fig. S2c**) and eliminated the response to ACh (**Fig. S2d**). Inserting mScarlet-I at the same location (B2) produced modest RET and partial responsivity, while fusing mScarlet-I to the C terminus (B3) produced no RET but recovered full responsivity (**Fig. S2c,d**). Fusions with mScarlet-I or mScarlet3 at the N-terminus of ACh-NeuBI0.4 displayed high RET (**Fig. S2c**) and robust responsivity (**Fig. S2d**). The mScarlet-I fusion demonstrated an even larger response than ACh-NeuBI0.4, and thus was designated Orange ACh-NeuBI0.5 and selected for further characterization.

To better characterize Orange ACh-NeuBI0.5’s response, we measured its spectral properties alongside those of its non-binding mutant, Orange ACh-NeuBI0.5mut, across a range of ACh concentrations. Interestingly, with increasing ACh concentration, ACh-NeuBI0.5 displayed an increase in not just total photonic output (**Fig. S3a**) but also in RET efficiency (**Fig. S3b**), whereas ACh-NeuBI0.5mut showed no significant change in either its RET efficiency or brightness following ACh treatment. The ACh-induced RET suggests a shift to an ACh-stabilized conformation with improved donor quantum yield, donor-acceptor distance, or donor-acceptor orientation. As a result, emission above 610 nm responds more strongly to ACh than overall emission (**Fig. S3c**), with 100 μM ACh eliciting a ∼6-fold signal increase. Its high responsiveness at the orange-red wavelengths, which are favorable for signal detection in tissue, makes it particularly suitable for in vivo applications.

We next characterized Orange ACh-NeuBI0.5 in HEK293A cells. By using the PDGFR transmembrane domain, Orange ACh-NeuBI0.5 was successfully anchored to the cell membrane and demonstrated consistent membrane targeting within HEK293A cells (**Fig. S4a**). Moreover, we observed a dose-dependent response to ACh treatment, with an up to 8-fold increase in luminescent signals (**Fig. S4b**).

To determine the specificity of Orange ACh-NeuBI0.5, we measured its K_d_ against a panel of neurotransmitters. Orange ACh-NeuBI0.5 binds to ACh with an apparent K_d_ of 898 μM in HEK293A cells, with a notably weaker affinity to choline (apparent K_d_ = 11 mM) (**Fig. S4c**). Orange ACh-NeuBI0.5 shows no detectable response to other neurotransmitters, including glutamate, GABA, serotonin, dopamine, glycine, and norepinephrine, suggesting specificity for ACh (**Fig. S4c**). Purified ACh-NeuBI0.5 protein exhibited a dose-dependent response to ACh treatment, with a K_d_ of 587 μM and a maximum response of ∼10-fold (**Fig. S4d**).

### Affinity maturation by deep mutagenesis via MIDAS

We next aimed to further improve the responsivity of Orange ACh-NeuBI0.5 to low concentrations of ACh. Even though ACh is estimated to transiently reach concentrations as high as 1–10 mM at the synaptic clefts of neuromuscular junctions^36^ and the release sites of striatal cholinergic interneurons^37,38^, Orange ACh-NeuBI0.5’s K_d_ of 898 μM is still far above the basal ACh concentrations in the brain (approximately 500 nM)^39^. We thus speculated that an ACh-NeuBI with higher affinity would be better suited for detecting changes in cholinergic signaling in vivo. Therefore, we performed a deep mutagenesis screening on Orange ACh-NeuBI0.5 to enhance its responsivity and affinity to enable robust detection of ACh in physiologically relevant low-concentration regimes.

We first used computational protein structural predictions to identify sites where mutations are predicted to improve protein folding in the ACh-bound state. We selected 16 sites in direct contact with ACh, in the second shell, at the interface between OpuBC and NanoBiT, and at the linker region (**Fig. 2a**). We then used MIDAS to perform enumerated saturation mutagenesis at each site, testing 4 concentrations of ACh in parallel in the primary screen (**Fig. 2b, Fig. S5a**). Mutations producing the largest responses to ACh were clustered at three second-shell locations, Gly-550, Leu-560, and Ala-610 (**Fig. 2b, Fig. S5a**). Specifically, G550Q, G550R, L560C, L560E, A610D, and A610E demonstrated up to 5-fold enhancements in K_d_ compared to Orange ACh-NeuBI0.5 (**Fig. 2c, Fig. S5b**). We then used MIDAS to improve Orange ACh-NeuBI0.5 further via saturation mutagenesis. As only Gln and Arg were beneficial at Gly-550, and this site was distant from the other two hotspots at position 560 and 610, we decided to fix position 550 as either Gln or Arg in the next round. We were concerned that the positively charged Arg may create electrostatic repulsion for ACh in some sectors of sequence and conformation space, so we selected to fix position 550 to Gln. Considering the potential for interactions between residues at positions Leu-560 and Ala-610 due to their spatial proximity, we then conducted enumerated saturated combinatorial mutagenesis at these positions. Several mutation combinations, designated QAD, QAE, QCD, QCE, QED, and QEE according to the amino acids at positions 550, 560, and 610, exhibited up to 2-fold and 3-fold improvements over Orange ACh-NeuBI0.5 in response to 10 and 100 µM ACh, respectively (**Fig. 2d, Fig. S6a**), improving in responsivity over G550Q alone. Variant QAE exhibited the highest affinity (K_d_ = 95 µM) and highest brightness at 10–100 µM ACh (**Fig. 2e right, Fig. S6b**), while QED exhibited decreased brightness in the absence of ACh (**Fig. 2e right, Fig. S6b**) and thereby the largest contrast between 0 and 10 or 100 uM ACh (**Fig. 2e left**). QED exhibited a 640% fluorescence increase to 100 μM ACh, representing 29-fold higher responsivity compared to the pre-MIDAS reporter NeuBI0.1.

**Fig. 2.**
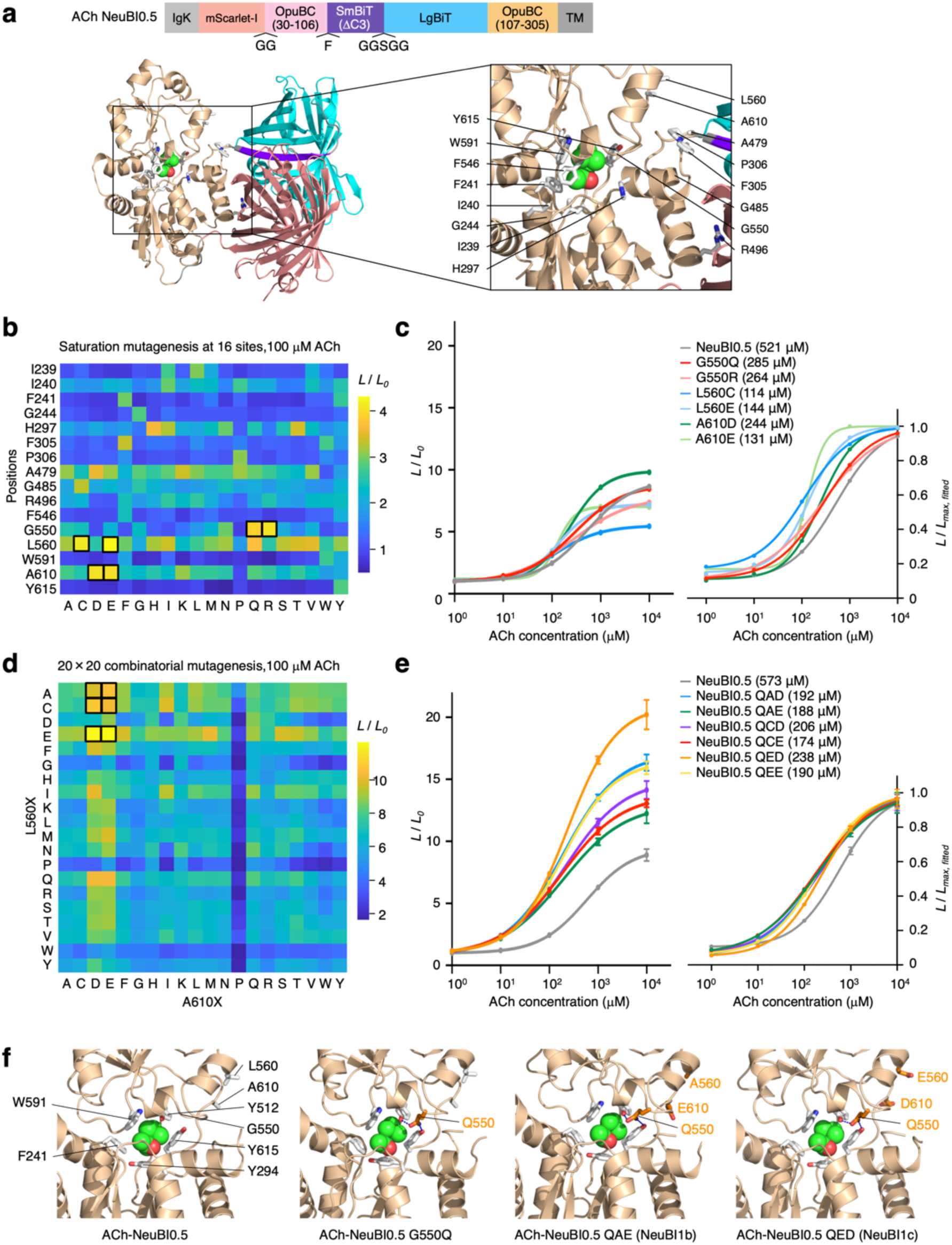
MIDAS applied for deep mutagenesis. **a**, Structure of ACh-NeuBI0.5 predicted by Chai-1. Wheat, OpuBC; salmon, mScarlet-I; cyan, LgBiT; purple, SmBiT; gray, linkers; green spheres, ACh. The side chains of the 16 residues selected for saturation mutagenesis are shown as white sticks. **b**, Heatmap of saturation mutagenesis at 16 sites by MIDAS, tested at 100 μM ACh, in HEK293A cells. Boxed cells indicate the top-performing mutants selected for further characterization. **c**, Dose-response curves of the plasmids encoding top mutants from the 16-site saturation mutagenesis in HEK293A cells, normalized to the H_2_O control (left), or normalized to fitted maximum (right). **d**, Heatmap of 20 × 20 combinatorial MIDAS at positions 560 and 610, tested at 100 μM ACh, in HEK293A cells. Boxed cells indicate the top-performing mutants selected for further characterization. **e**, Dose-response curves of the plasmids of top mutants from the 20 × 20 combinatorial mutagenesis in HEK293A cells, normalized to the H_2_O control (left), or normalized to fitted maximum (right). **c, e**, Curves were fitted using the “nonlinear regression (log(inhibitor) vs. response – variable slope)” model in Prism. **f**, Structures of NeuBI0.5, NeuBI0.5 G550Q, NeuBI1b, and NeuBI1c predicted by Chai-1. Wheat, OpuBC; green spheres, ACh. The side chains of residues in direct contact of ACh are shown as white sticks. The side chains of mutated residues are shown as orange sticks.

We sought to understand the physical basis for the stepwise improvement in affinity from ACh-NeuBI0.5 to NeuBI.05 G550Q and NeuBI0.5 QAE or QED. We used the AlphaFold3 algorithm^40^, accessed via the public Chai-1 server^41^, to predict the structure of each variant bound to ACh. Chai-1 accurately recapitulated the known OpuBC-ACh cocrystal structure (PDB: 6V1R) and predicted no significant changes in the ACh binding pocket for an OpuBC domain with the NeuBI-ACh0.5 sequence (**Fig. 2f**). In ACh-NeuBI0.5 G550Q, the Gln sidechain at position 550 was predicted to engage in hydrogen bonds with Tyr-512 and Tyr-615 which are situated on the two lobes on opposite sides of the ACh-binding pocket. Thus Gln-550 may help stabilize the closed conformation of the OpuBC domain around the bound ACh.

The effects of mutations at 560 and 610 are less obvious from the structural predictions, as the conformations of side chains forming the ACh-binding pocket are not predicted to change. As QAE exhibits higher affinity than G550Q alone, L560A and E610D may subtly stabilize the closed conformation of the OpuBC domain. As these side chains face solvent, the effects of the mutations may be to optimize the water shell in their vicinity in the closed state. In contrast, the benefit of QED in responsivity derives primarily from lowering baseline bioluminescence rather than improving affinity relative to G550Q alone, so the combination of L560E and A610D apparently destabilizes SmBiT-LgBiT complementation in the absence of ACh. As these side chains are located close to SmBiT in the AlphaFold3-predicted structure, it is possible they have a gate-keeping function against spontaneous SmBiT-LgBiT complementation.

### Responsivity, specificity, and reversibility of optimized ACh-NeuBIs

We designated ACh-NeuBI0.5 QAE as ACh-NeuBI1b for its highest brightness at all ACh concentrations among all variants, and ACh-NeuBI0.5 QED as ACh-NeuBI1c for its highest contrast, and tested their performance in cells and in vitro. Both proteins expressed well at the plasma membrane (**Fig. S7a**) and maintained selectivity for ACh over choline (**Fig. S7b**). Additionally, their RET efficiencies both improved with increasing ACh concentrations (**Fig. S7c)**, resulting in higher responses at wavelengths above 610 nm (**Fig. S7d**). This feature makes them especially well-suited for in vivo applications. We next tested the characteristics of NeuBI1b and NeuBI1c in free soluble form. Purified untethered NeuBI1b and NeuBI1c proteins were enhanced in terms of their ACh responsivity and selectivity over choline (**Fig. S7e, Fig. S4d**), but surprisingly did not exhibit higher affinity to ACh compared to NeuBI0.5 in vitro (**Fig. S7e, Fig. S4d**). This suggests that our NeuBI improvements which were identified by screening on mammalian cell surfaces may not have been successfully identified if screened in a different context, e.g. as soluble bacterial periplasmic proteins.

To explore ACh-NeuBI applicability in more physiologically relevant settings, we extended our analysis to primary cortical neurons. Remarkably, ACh-NeuBIs displayed excellent membrane targeting, across cell bodies, dendrites, and spines (**Fig. S8a**). NeuBI0.5 exhibited up to a 60-fold increase in luminescence in an ACh concentration-dependent manner (**Fig. S8b**), while NeuBI1b and NeuBI1c demonstrated even greater responses, achieving up to a 200-fold increase (**Fig. S8c-d**). Notably, their ACh responsivity exceeded that observed in HEK293A cells (**Fig. 2e**, **Fig. S8b-d**), highlighting the enhanced responsivity of these sensors in the native neuronal environment, which is likely due to improved trafficking and enhanced membrane localization. Moreover, all NeuBIs displayed exceptional selectivity for ACh over choline in primary cortical neurons (**Fig. S8b-d**). These findings suggest that ACh-NeuBIs are well-suited for studying cholinergic transmission under physiologically relevant conditions.

### Optimized ACh-NeuBIs robustly respond to ACh in living mice

To investigate ACh-NeuBIs’s responsiveness to ACh in mice, we expressed NeuBI0.5, NeuBI1b, or NeuBI1c in nude mice by hydrodynamic transfection. After 24 hours, the mice were intraperitoneally (i.p.) injected with NanoLuc substrate, fluorofurimazine (FFz)^42^, and were imaged for ∼3 min until signals reached a plateau. Once the plateau was reached, mice were administered saline or ACh at various concentrations, and imaging was resumed immediately (**Fig. 3a**). In NeuBI0.5-expressing mice, ACh administration at 0.1 mg/kg, 1 mg/kg, and 10 mg/kg resulted in significant signal increases of 3.5-fold, 7.5-fold, and 30-fold compared to saline, respectively, demonstrating the robust responsiveness of ACh-NeuBI0.5 in mice (**Fig. 3b**,**c**). Compared to NeuBI0.5, NeuBI1b and NeuBI1c demonstrated further improvements at all tested amounts of ACh. Specifically, NeuBI1b and NeuBI1c exhibited signal increases of 10.6- and 10.2-fold at 0.1 mg/kg, 38- and 35-fold at 1 mg/kg, and 103- and 173-fold at 10 mg/kg (**Fig. 3b**,**c**). Thus, NeuBI1b and NeuBI1c detect ACh larger signal changes than NeuBI0.5 across the entire concentration range in vivo, confirming earlier in vitro results. These results thus demonstrate the utility of MIDAS in discovering protein optimizations that are useful in vivo, as well as in vitro.

**Fig. 3.**
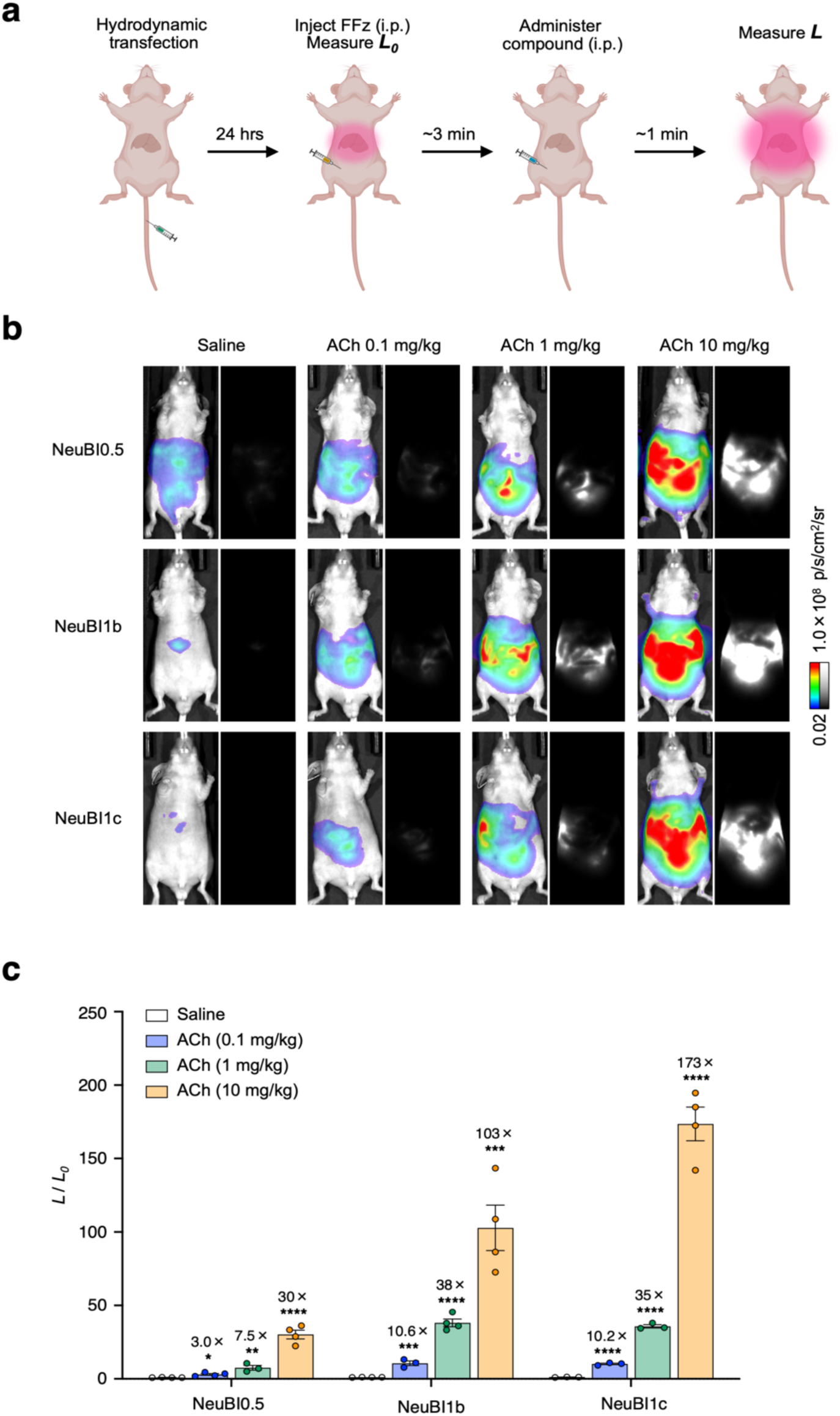
In vivo validation of NeuBIs. **a**, Experimental scheme for testing NeuBI testing in the liver. NeuBI plasmids are hydrodynamically transfected into nude mice. After 24 h, FFz is injected intraperitoneally (i.p.) to measure baseline luminescence. Once a plateau is reached, saline or ACh is injected i.p., and imaging is continued. **b**, Representative bioluminescence images acquired after the treatment of saline or ACh (0.1 mg/kg, 1 mg/kg, or 10 mg/kg) in NeuBI0.5-, NeuBI1b-, or NeuBI1c-expressing mice. **c**, Fold of signal increase in response to ACh (0.1 mg/kg, 1 mg/kg, or 10 mg/kg), normalized to the saline control. ns, not significant; *, p < 0.05; **, p < 0.01; ***, p < 0.001; **** p < 0.0001, by unpaired two-tailed Student’s t-test compared to the saline control. Error bars, SEMs.

To rule out the possibility of ACh influencing substrate delivery or other non-specific factors unrelated to ACh binding to NeuBIs, we hydrodynamically transfected a mScarlet-I-NanoLuc plasmid, which is unregulated by ACh, into nude mice. Subsequent measurements of their responses to ACh and saline revealed no significant differences between the saline and ACh treatment groups (**Fig. S9**). This finding indicates that the observed response was specifically triggered by ACh binding to NeuBIs.

## Discussion

In this study, we engineered ACh Neurotransmitter Bioluminescent Indicator (ACh-NeuBI) for real-time monitoring of ACh dynamics in vivo. ACh-NeuBI was engineered by integrating a cpNanoLuc into optimized ACh-binding OpuBC domains, with subsequent optimization using an efficient high-throughput strategy we refer to as microbe-independent deep assembly and screening (MIDAS). The final optimized variants, ACh-NeuBI1b and ACh-NeuBI1c, demonstrate exceptional responsivity, specificity, and reversibility, enabling robust detection of ACh dynamics in both primary neurons and living mice.

The MIDAS strategy we developed provides a rapid, high-throughput approach for optimizing protein function directly in mammalian cells, bypassing traditional microbial cloning steps. Traditionally, plasmid cloning has been a necessary step for site-directed mutagenesis or linker variation in protein engineering in mammalian cells. However, cloning mutant genes is a complex, multi-step, and lengthy process of in vitro PCR, vector preparation, plasmid assembly or ligation, transformation, colony picking, bacterial culture, and plasmid purification, thus requiring substantial resources, labor and time. Plasmid purification alone is a multi-step process requiring numerous reagents, supplies, and careful manual procedures. In contrast, MIDAS bypasses the entire cloning procedure, requiring simply two PCR reactions followed immediately by transfection, thereby reducing the time between primer receipt and transfection to one day.

Using this streamlined MIDAS approach, we rapidly improved the output dynamic range of ACh-NeuBI from 20% to ∼200% (3-fold) at 100 μM ACh in just three rounds, then further enhanced it to 8-fold with two additional rounds of screening. Importantly, the molecular changes in our final round of optimization do not contact ACh and are not predicted to alter protein folding, but may instead affect weak interactions between domains or between side chains and solvent. As current computational algorithms based on machine learning of mostly single-domain solvent-free protein structures do not predict such interactions well, these types of improvements must be obtained through empirical screening, which MIDAS greatly facilitates.

MIDAS is particularly well suited for saturation mutagenesis at targeted sites, either at a series of individual sites independently or at multiple sites simultaneously. We note that targeted mutagenesis is required for fully sampling sequence space at any given site. Random mutagenesis is useful to identify sites where mutations can improve protein function, but is incomplete in coverage at each codon. In one study, a mutagenesis rate of 1.7% maintained functionality in 6.7% of sequences constructed by error-prone PCR^43^. At this rate, the frequency of any codon receiving more than one mutation while appearing within the functional subpopulation is 0.0172 × 0.067, or 1 in 50,000. Likewise, a mutagenesis rate of 10% maintained functionality in 0.17% of sequences, yielding a recoverable double-mutation rate per codon of 0.12 × 0.0017, or 1 in 59,000. Thus increasing mutagenesis rates to improve double-mutation rates reduces recoverability in a corresponding manner, such that in practice the isolation of variants with two useful mutations per codon is excessively rare from randomly generated libraries. Thus, practically speaking, RM can be considered to cover only the ≤9 amino acid changes possible from single mutations in a codon. Increasing mutagenesis depth at hotspots to optimize protein function thus requires targeted saturation mutagenesis at interesting sites. Mutant libraries of all possible amino acid combinations at P positions (e.g. encoded as NNS or NNK) can be expressed and screened in 32^P^ × 10 cells, where the oversampling factor of 10 is required to reach 99% probability that all possible combinations are present, given Poisson noise in sampling^44^. Screening of targeted saturation mutagenesis libraries in mammalian is straightforward at one site, requiring screening only 20 enumerated clones or 320 samples of a NNS/NNK library, but becomes cumbersome to scale to multiple sites with earlier methods, with just two sites requiring the generation of 400 clones deterministically or the screening of >10000 NNS/NNK libraries. By allowing deterministic construction and characterization directly in mammalian cells, MIDAS avoids the need to clone each combination in bacteria and prepare plasmid DNA individually, or to screen in bacteria and validate in mammalian cells, thereby saving substantial time and expense.

In addition to accelerating sensor optimization, MIDAS enables the rapid generation of large, high-quality datasets for training machine learning models on enzyme or binder structure-function relationships. A major challenge in applying machine learning to predict the effects of mutations on dynamic aspects of protein function, such as enzyme catalysis or conformational changes, is the limited availability of high-quality experimental datasets, particularly for functional optimization in mammalian systems. MIDAS addressed this challenge by enabling the systematic and rapid screening of mutants directly in mammalian cells, thus providing rich and reliable experimental data that can refine machine learning algorithms for rational protein engineering.

Compared to existing fluorescent ACh indicators, such as GRAB-ACh3.0^24^ and iAChSnFR^5^, ACh-NeuBI1b and ACh-NeuBI1c offer advantages in input dynamic range, output dynamic range, and lack of biological perturbation. GRAB-ACh3.0 and iAChSnFR showed approximately 3-fold and 8-fold responses, respectively, to 100 µM ACh treatment in cultured HEK cells, with their signals peaking at this concentration. ACh-NeuBI1b/c responses were comparable to iAChSnFR at 100 µM ACh, but continued to increase with higher ACh concentrations to reach 15- to 20-fold maximum changes. Thus, the higher K_d_ of ACh-NeuBI1b/c and their larger overall responsivity extends their input dynamic range without sacrificing responses at intermediate inputs, and achieves responses across a broader range of ligand concentrations without reaching saturation at physiological concentrations. This will be especially useful in locations where ACh concentrations can exceed 100 µM, such as the synaptic clefts of neuromuscular junctions and cholinergic interneuron release sites in the striatum^36–38^. Furthermore, a high K_d_ reduces competition with endogenous ACh receptors, ensuring that the sensor minimally perturbs natural neuronal activity while still accurately reporting fluctuations in ACh levels. This is desirable in locations with low-affinity ACh receptors, such as skeletal muscles which express α7 nAChRs with K_d_ ∼180 µM^45^.

In contrast to fluorescent ACh indicators, ACh-NeuBI offers a noninvasive method for monitoring ACh dynamics in vivo. Imaging of fluorescent indicators in deep locations requires the implantation of optical elements such as lenses or fibers for excitation light delivery and emission light collection. These can damage tissues and compromise animal health, potentially leading to artificial or unreliable results. Moreover, the complex surgeries and imaging setups required for fluorescent indicators demand extensive training and effort. Stable implantation of optical elements may be impossible when the imaging location is within or underneath motile tissues, an example being the liver and other organs of the gastrointestinal tract which are located beneath the motile abdominal wall. In contrast, the bioluminescent ACh-NeuBI enables noninvasive and convenient imaging with cameras outside the body in an inexpensive lightproof box or dark room. The absence of excitation light and autofluorescence significantly reduces background noise, thereby enhancing the detection of changes in photonic emissions from deep tissues without requiring implanted optical components. Furthermore, the noninvasive nature of ACh-NeuBI enables its application in peripheral systems and minimizes disturbances caused by invasive techniques, thereby maintaining the natural integrity of the biological systems being studied.

The modular architecture of ACh-NeuBI can serve as a template for generating other NeuBIs. By substituting the ACh-binding OpuBC domains with binding domains specific to another neurotransmitter, and employing a systematic design approach – including initial screening of configurations followed by MIDAS-based optimization of linkers and binding affinities – this versatile framework can be customized to monitor any neurotransmitter of interest. Consequently, ACh-NeuBI should be a generalizable design that can be adapted to diverse neurotransmitters to allow noninvasive exploration of their distributions in vivo.

The inherently genetically encoded nature of ACh-NeuBI enables targeted investigation of cholinergic activity within specific cell populations by employing cell type-specific promoters. This approach is particularly advantageous in heterogeneous brain tissues, as it allows the isolation and detailed study of distinct cell types, such as various types of neurons and glial cells. By targeting these specific populations, ACh-NeuBI could provide valuable insights into complex neurological functions and disorders, including memory^17^, attention^18^, pain^46^, and neuroinflammation^47^.

In summary, we developed ACh-NeuBI, a genetically encoded bioluminescent indicator for real-time, noninvasive imaging of cholinergic signaling in vivo. To rapidly optimize ACh-NeuBI, we developed MIDAS, a high-throughput protein engineering platform that bypasses traditional cloning. By systematically refining interdomain linkers and mutational hotspots, we successfully enhanced the responsivity, specificity, and dynamic range of ACh-NeuBI within several rounds of mutagenesis. The final optimized variants, ACh-NeuBI1b and ACh-NeuBI1c, exhibited superior signal contrast, enabling robust detection of ACh dynamics in primary neurons and living mice, highlighting the power of MIDAS for accelerating protein optimization in mammalian systems.

## Supplementary Information

**Fig. S1.**
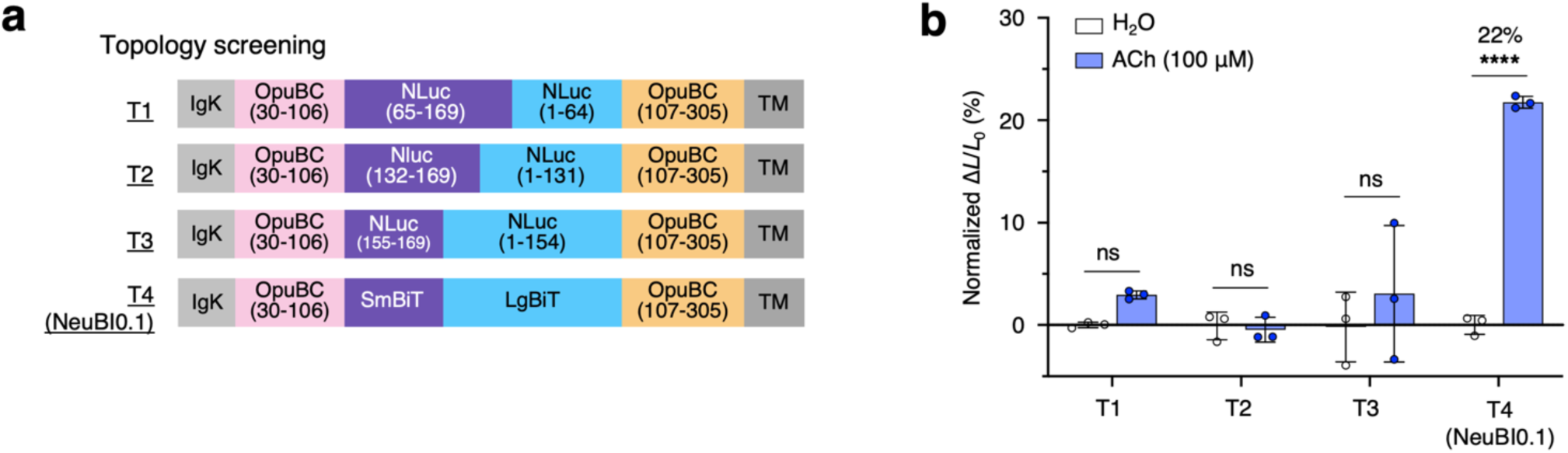
Topology screening. **a**, Topologies T1–T4 testing various circular permutation sites of NanoLuc. **b**, luminescent signal change of T1–T4 upon the treatment of ACh (100 μM), normalized to the H_2_O control. *, p < 0.05; **, p < 0.01; ***, p < 0.001; **** p < 0.0001, by unpaired two-tailed Student’s t-test compared to the H_2_O control. Error bars, SEMs. Curve fitted by the “nonlinear regression (log(inhibitor) vs. response – variable slope)” model in Prism.

**Fig. S2.**
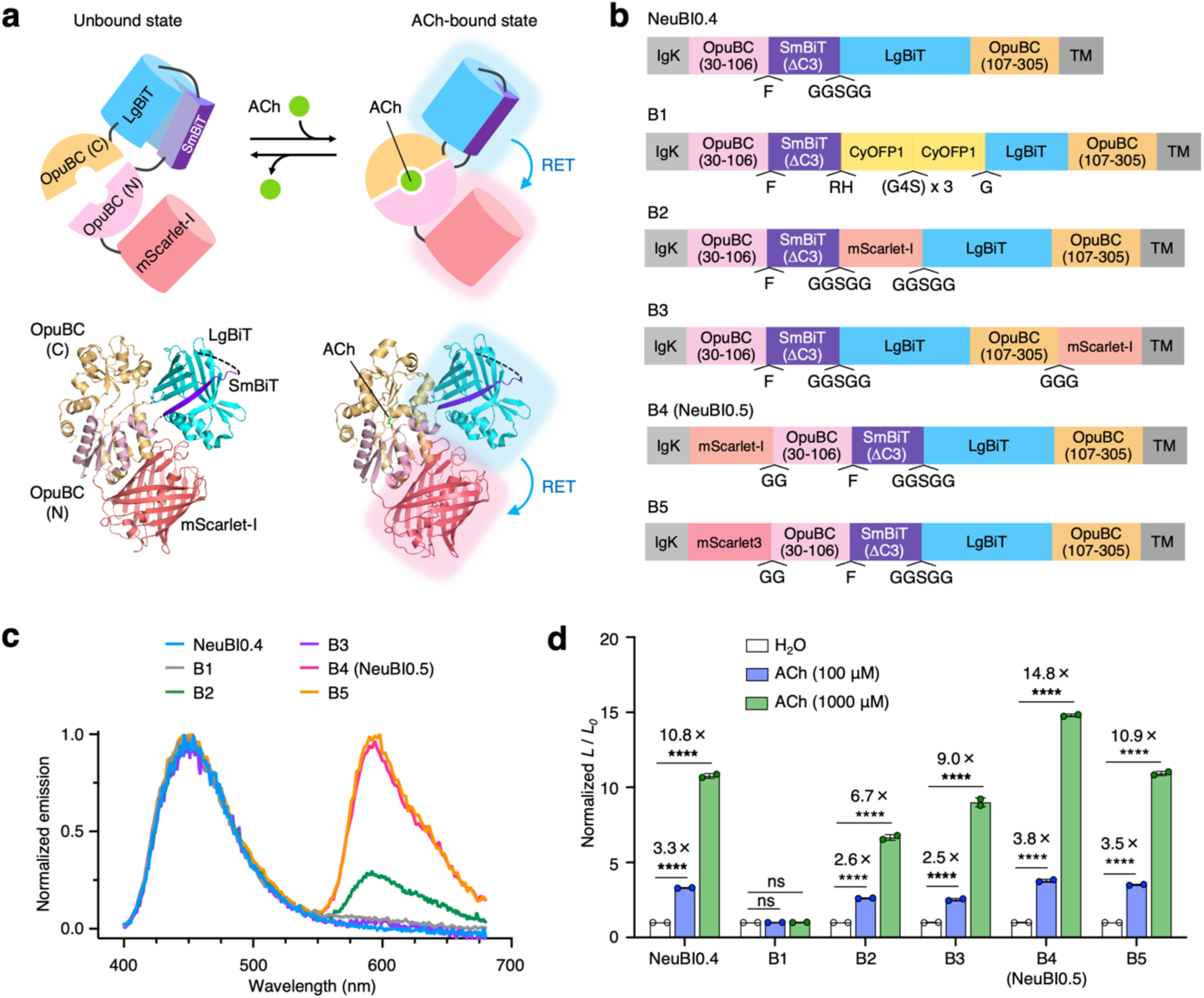
Engineering an orange ACh-NeuBI0.5. **a**, Proposed mechanism of Orange ACh-NeuBI0.5. Top, cartoon scheme of Orange ACh-NeuBI0.5. Bottom, the structure model of Orange ACh-NeuBI0.5, modified from PDB files 6URU (OpuBC), 5IBO (NanoLuc), and 5LK4 (mScarlet). ACh binding to OpuBC leads to an increase in the luminescence of cpNanoLuc and red light emission from mScarlet-I via resonance energy transfer (RET). **b**, Topologies of constructs B1–B5 for red-shifting spectra. **c**, Spectra of B1–B5, measured in the presence of ACh (1000 μM). **d**, Fold of signal increase of B1–B5 in response to 100 μM ACh or 1000 μM ACh, normalized to the H2O control. ns, not significant; *, p < 0.05; **, p < 0.01; ***, p < 0.001; **** p < 0.0001, by unpaired two-tailed Student’s t-test compared to the H_2_O control. Error bars, SDs.

**Fig. S3.**
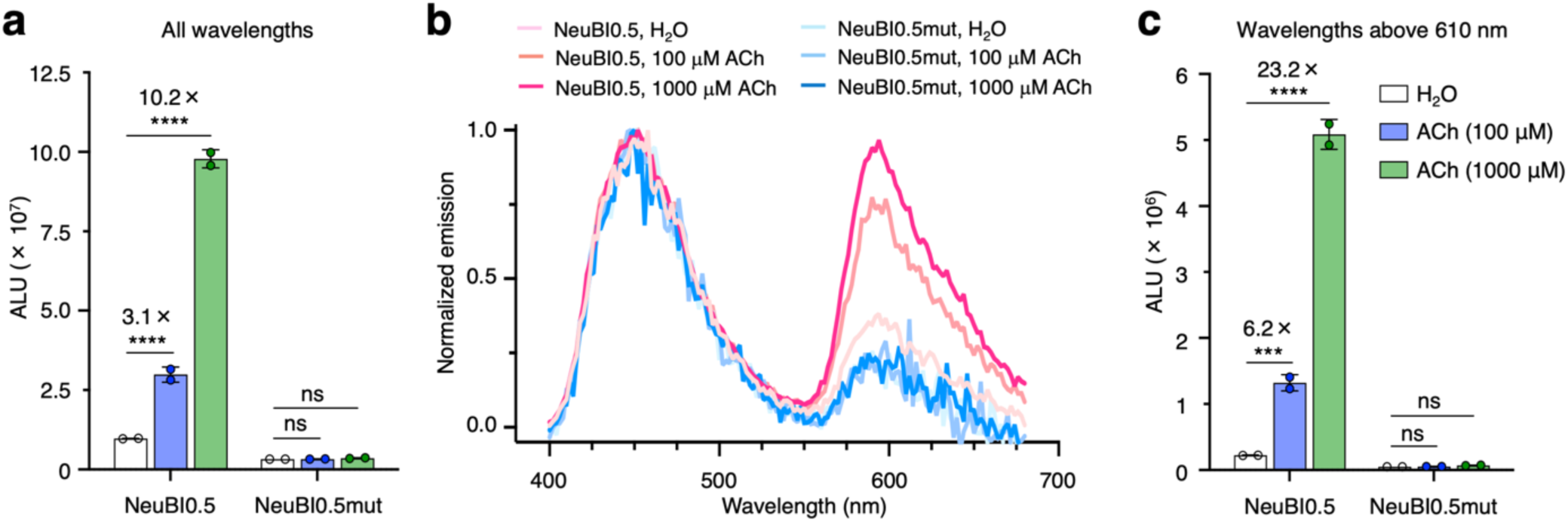
Optical components of the response of ACh-NeuBI0.5. **a**. Luminescence of ACh-NeuBI0.5 and ACh-NeuBI0.5mut measured without filter, without ACh, with 100 μM ACh, or with 1000 μM ACh. ALU, arbitrary luminescence units. **b**, Spectra of ACh-NeuBI0.5 and ACh-NeuBI0.5mut, measured without ACh, with 100 μM ACh, or with 1000 μM ACh. **c**. Luminescence of ACh-NeuBI0.2 and ACh-NeuBI0.2mut measured with a 610 nm longpass filter, without ACh, with 100 μM ACh, or with 1000 μM ACh. ALU, arbitrary luminescence units. **a,c**, ns, not significant; *, p < 0.05; **, p < 0.01; ***, p < 0.001; **** p < 0.0001, by unpaired two-tailed Student’s t-test compared to the H_2_O control. Error bars, SDs.

**Fig. S4.**
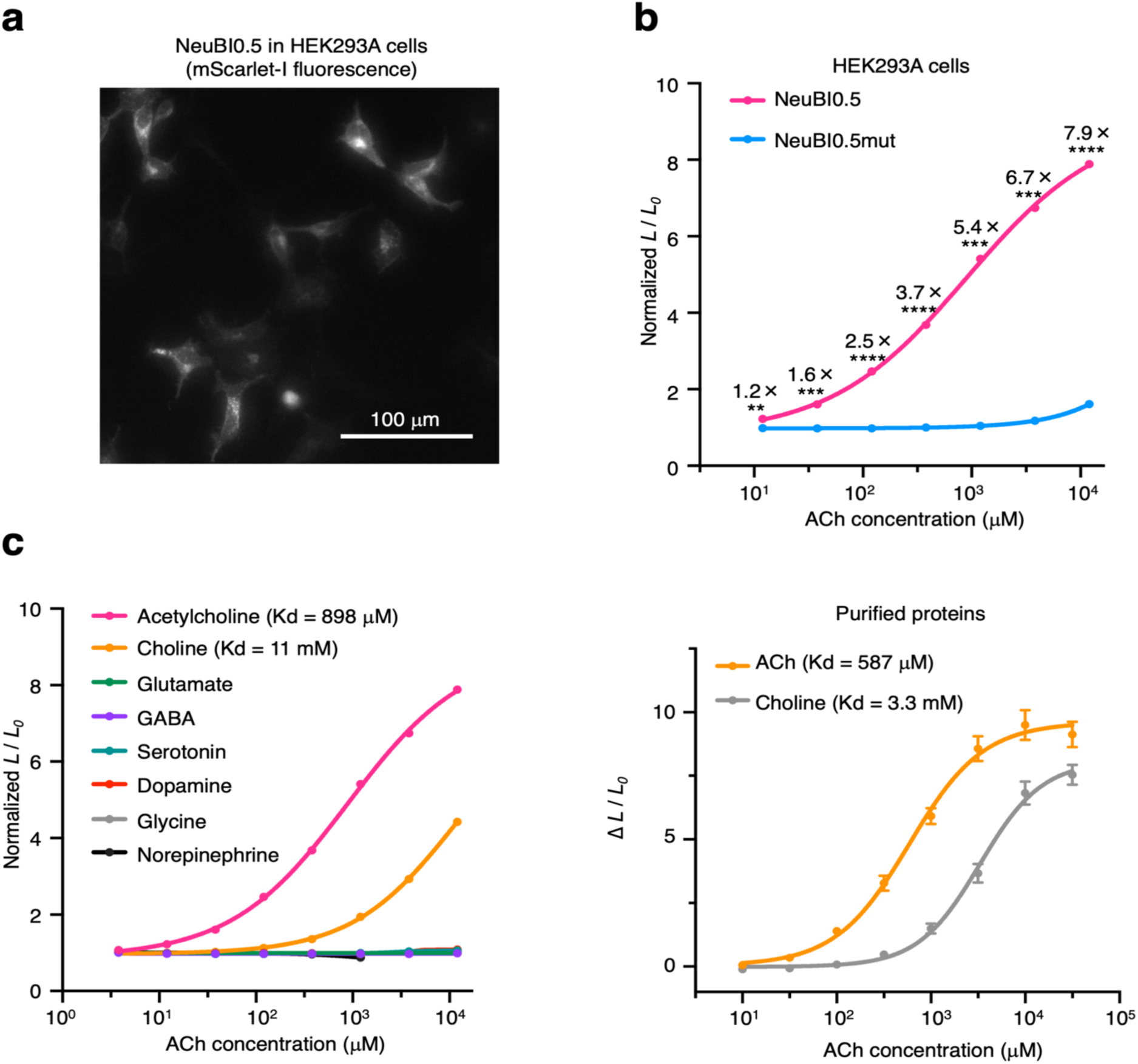
Characterization of ACh-NeuBI0.5. **a**, mScarlet-I fluorescence of Orange ACh-NeuBI0.5 in HEK293A cells. **b,** Fold of signal increase of Orange ACh-NeuBI0.5 in response to ACh at different concentrations in HEK293A cells, normalized to the H_2_O control. **, p < 0.01; ***, p < 0.001; **** p < 0.0001, by unpaired two-tailed Student’s t-test compared to the H_2_O control. Error bars, SEMs. **c**, Fold of signal increase of Orange ACh-NeuBI0.5 in response to a panel of neurotransmitters at different concentrations in HEK293A cells. Error bars, SEMs. **d**, Dose-dependent response of purified NeuBI0.5 proteins to ACh treatment. Error bars, SEMs. **b-d,** Curves fitted by the “nonlinear regression (log(inhibitor) vs. response – variable slope)” model in Prism.

**Fig. S5.**
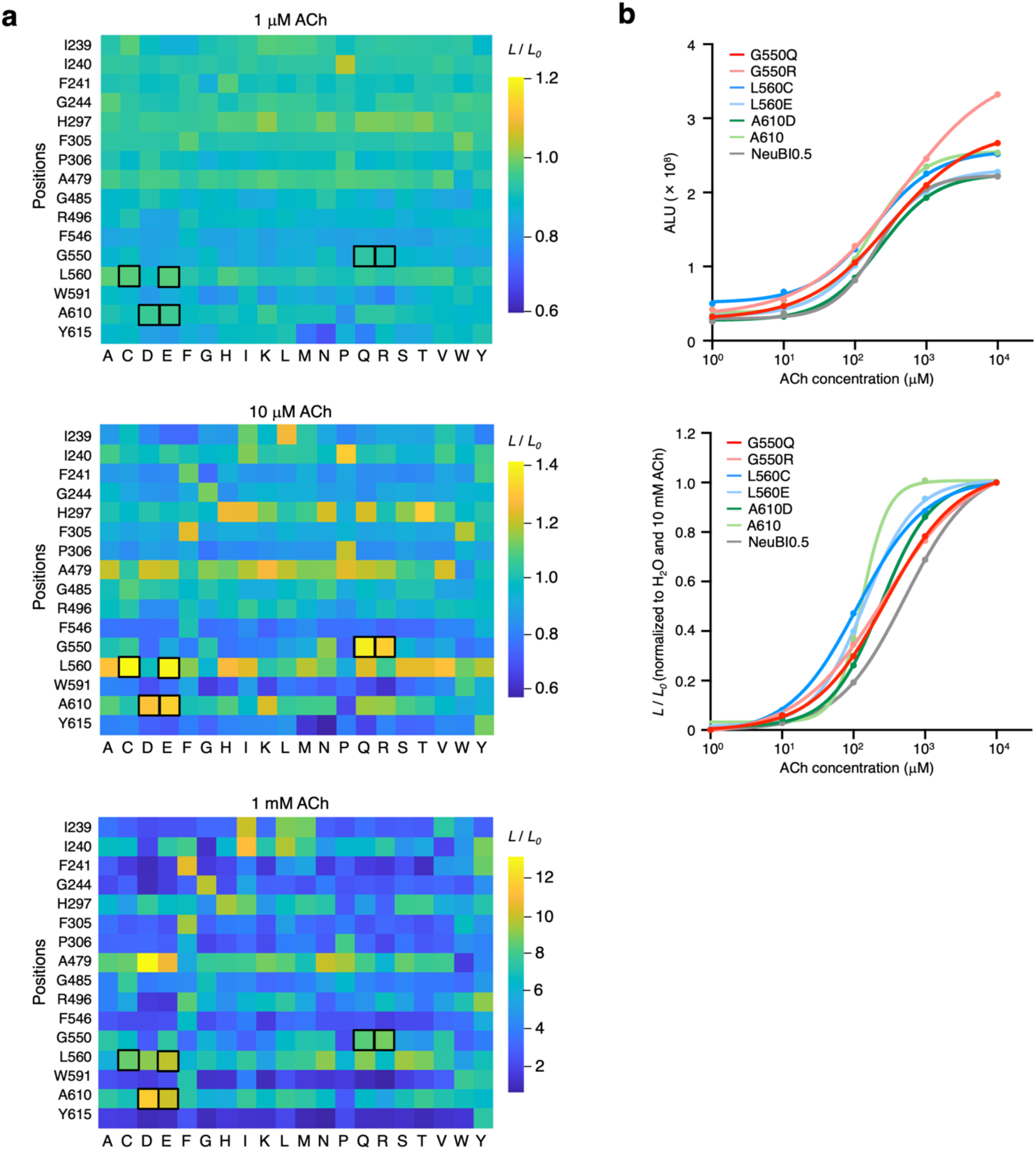
MIDAS results from saturation mutagenesis at 16 sites. **a**, Heatmap showing MIDAS results from saturation mutagenesis at 16 sites in HEK293A cells, tested at 1 μM (top), 10 μM (middle), and 1 mM (bottom) ACh. Boxed cells indicate the top-performing mutants selected for further characterization. **b**, Dose-response curves of the top mutants from the 16-site saturation mutagenesis in HEK293A cells, raw signals (top), or responses normalized to H_2_O and 10 mM ACh (bottom). All curves were fitted using the “nonlinear regression (log(inhibitor) vs. response – variable slope)” model in Prism.

**Fig. S6.**
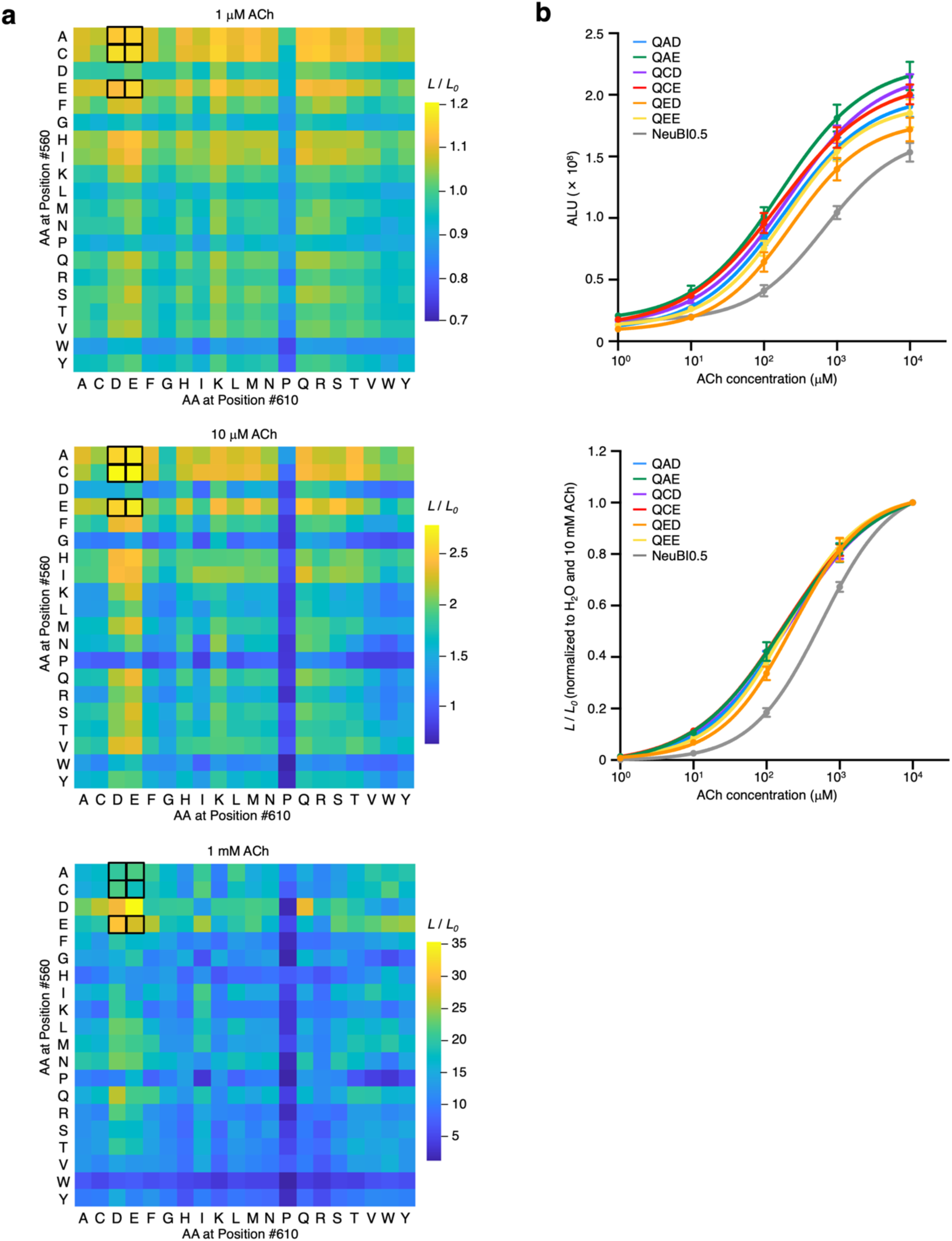
MIDAS results from 20 × 20 combinatorial mutagenesis at positions 560 and 610. **a**, Heatmap showing MIDAS results from 20 × 20 combinatorial mutagenesis at positions 560 and 610 in HEK293A cells, tested at 1 μM (top), 10 μM (middle), and 1 mM (bottom) ACh. Boxed cells indicate the top-performing mutants selected for further characterization. **b,** Dose-response curves of the top mutants from the 20 × 20 combinatorial mutagenesis in HEK293A cells, raw signals (top), or responses normalized to H_2_O and 10 mM ACh (bottom). All curves were fitted using the “nonlinear regression (log(inhibitor) vs. response – variable slope)” model in Prism.

**Fig. S7.**
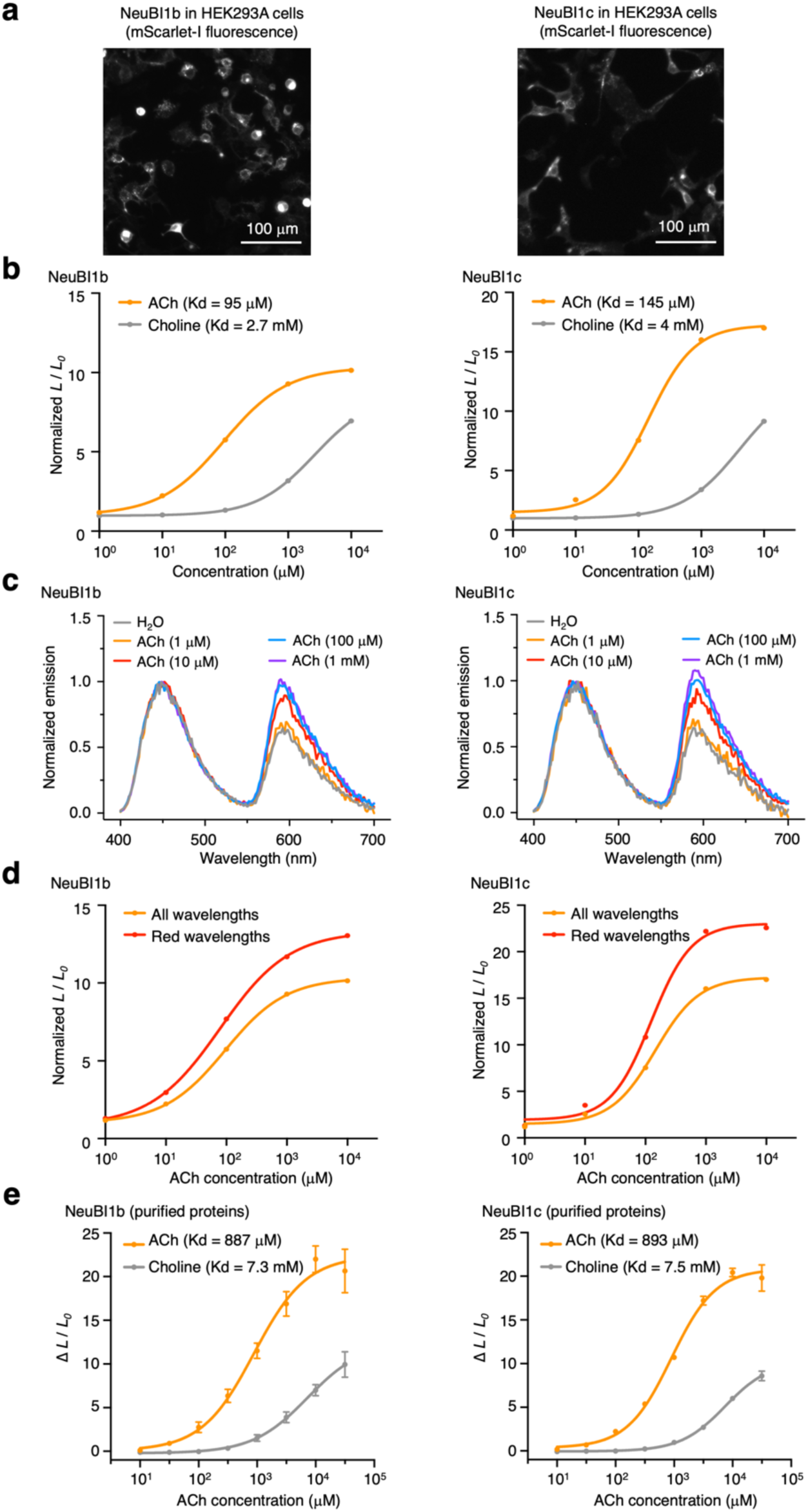
Characterization of ACh-NeuBI1b and NeuBI1c. **a,** mScarlet-I fluorescence of NeuBI1b (left) and NeuBI1c (right) in HEK293A cells. **b,** Fold of signal increase of NeuBI1b (left) and NeuBI1c (right) in response to ACh or choline at different concentrations in HEK293A cells, normalized to the H_2_O control. **c**, Spectra of NeuBI1b (left) and NeuBI1c (right), measured without ACh, or with ACh at various concentrations. **d**, Fold of signal increase of NeuBI1b (left) and NeuBI1c (right) in response to ACh at different concentrations in HEK293A cells without filter (orange line) or with a 610 nm longpass filter (red line). **e**, Dose-dependent response of purified NeuBI1b (left) and NeuBI1c (right) proteins to ACh treatment. Error bars, SEMs. **b,d,e,** Curves fitted by the “nonlinear regression (log(inhibitor) vs. response – variable slope)” model in Prism.

**Fig. S8.**
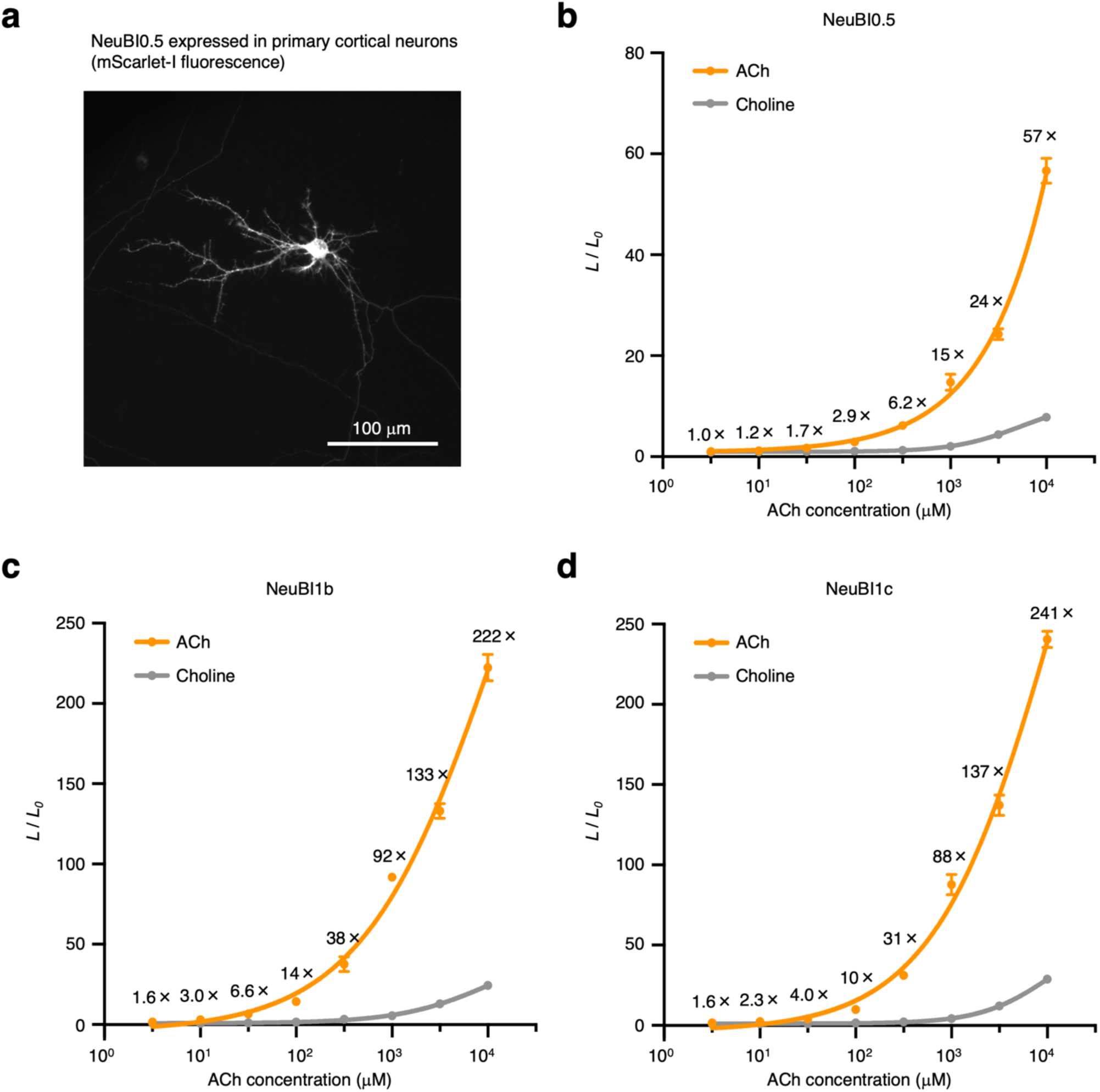
Characterization of ACh-NeuBIs in cultured neurons. **a**, mScarlet-I fluorescence of ACh-NeuBI0.5 in primary cortical neurons. **b-d**, Fold of signal increase of NeuBI0.5 (**b**), NeuBI1b (**c**), and NeuBI1c (**d**) in response to ACh or choline at different concentrations in primary cortical neurons. Error bars, SEMs. Curves fitted by the “nonlinear regression (log(inhibitor) vs. response – variable slope)” model in Prism.

**Fig. S9.**
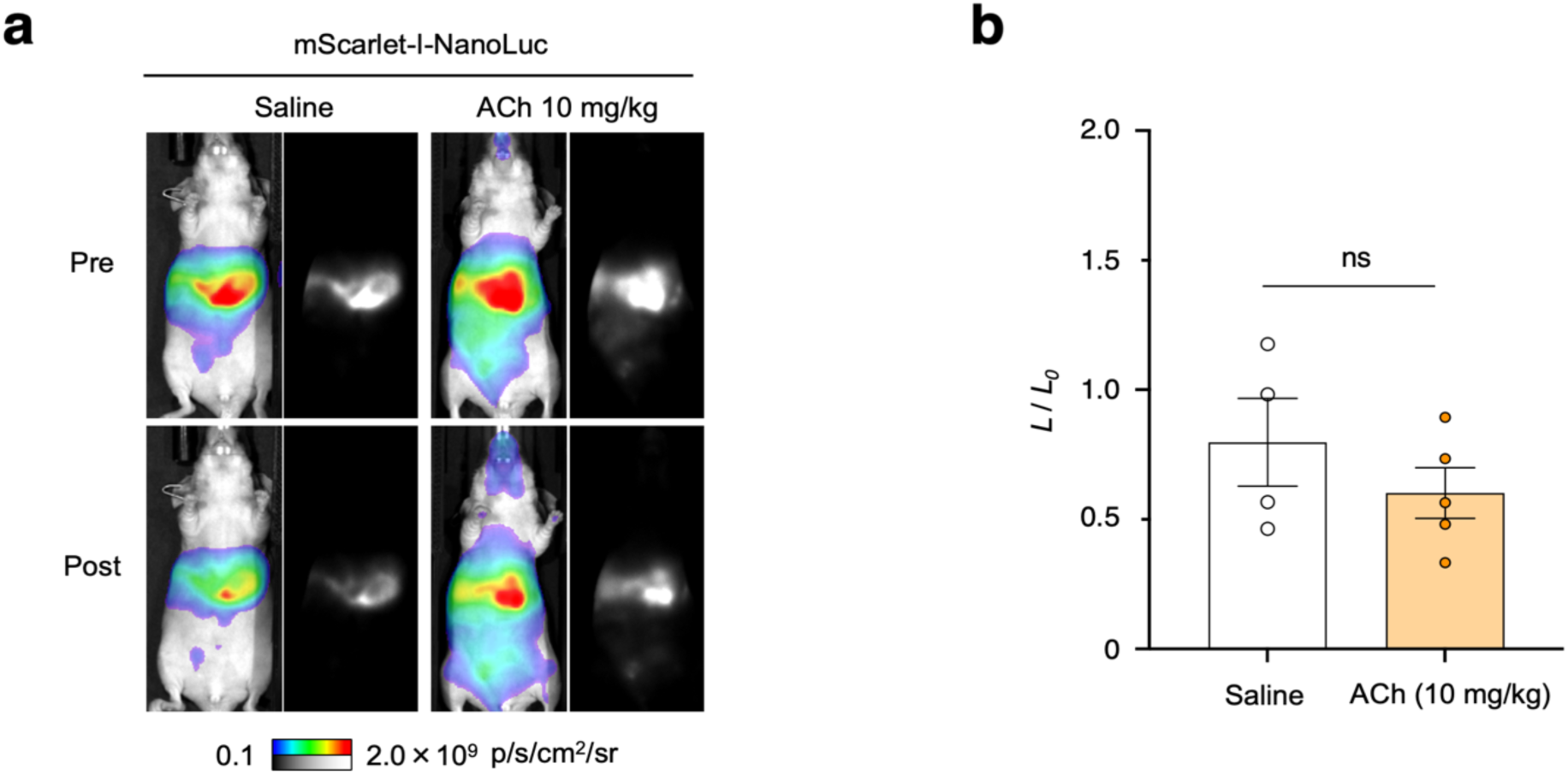
ACh doesn’t affect NanoLuc’s brightness in vivo. **a**, Representative bioluminescence images acquired before and after the treatment of saline or ACh (10 mg/kg) in mScarlet-I-NanoLuc-expressing mice. **b**, Fold of signal increase in response to ACh (10 mg/kg), normalized to the saline control. ns, not significant, by unpaired two-tailed Student’s t-test. Error bars, SEMs.

## Methods

### Chemicals

The following compounds were purchased: acetylcholine chloride, choline chloride, L-glutamate potassium monohydrate, serotonin hydrochloride, dopamine hydrochloride, and glycine (Sigma-Aldrich); (-)-norepinephrine (Selleck Chemicals); and GABA (Tocris).

### Molecular Cloning

DNA primers for molecular cloning were synthesized by Integrated DNA Technologies. Optimized OpuBC domains were PCR amplified from pAAV.CAG.iAChSnFR, a gift from Loren Looger (Addgene 137955)^5^. NanoLuc was PCR amplified from pF4Ag.NanoLuc (ATG-42), a gift from Lance Encell (Addgene 137777). mScarlet-I was PCR amplified from Lck-mScarlet-I, a gift from Dorus Gadella (Addgene 98821). PCR amplification was conducted with 20 to 40 bp primers with ∼20 bp overhangs and the Phusion Flash High-Fidelity PCR Master Mix (Thermo Scientific). Molecular cloning was carried out using In-Fusion HD cloning kit (Takara Bio) with a plasmid vector linearized by restriction enzymes and 1-2 DNA inserts with ∼20 bp overhangs. Sequences of constructed plasmids were verified using Sanger sequencing (Elim Biopharm) or whole-plasmid sequencing (Primordium Labs). The intermediate ACh-NeuBI constructs were assembled on the pc3 vector (Addgene) behind the CMV promoter. The best ACh-NeuBI indicator, ACh-NeuBI0.2, and ACh-NeuBI0.2mut were then cloned into the AAV packaging vector pAAV under the CMV promoter.

### MIDAS

The region of interest (ROI) for mutagenesis was identified within the sensor’s coding sequence, based on structural information. One pair of forward and reverse primers were designed to amplify the entire sequence, including the CMV promoter, the Kozak sequence, ACh-NeuBI’s coding sequence, and the bGH poly(A) termination sequence. Another set of primers were designed around the ROI introduce mutations. The reverse primers were designed to completely overlap with the template sequence right before the ROI, while the forward primers were designed to include an 18–24 bp overhang with the template before the ROI, followed by mutation-encoding sequences within the ROI, and ending with an 18–24 bp sequence that overlaps with the template after the ROI. For double-site combinatorial mutagenesis screening, two sets of primers were designed around the two ROIs following the same principles. All primers were designed to have a melting temperature of 60-66°C.

Primary PCR reactions used plasmid as template. The shared primary PCR fragments, which can be used as templates in multiple secondary PCR reactions, were gel-purified using a standard gel extraction protocol. The unique primary PCR fragments were verified by gel electrophoresis, then diluted 50-100× for use as templates in the secondary PCR. The secondary PCR reactions were conducted using gel-purified shared PCR fragments and diluted unique PCR fragments as templates, using F1 and R2 primers. All PCR reactions were conducted using Phusion Flash High-Fidelity PCR Master Mix (Thermo Scientific), following the manufacturer’s instructions.After verifying the integrity of the secondary PCR products using gel electrophoresis, 1 μL of the PCR products were transfected in HEK293A cells using lipofectamine 3000 (Life Technologies), following the manufacturer’s instructions. The mutants were characterized 24–48 h later through luminescence measurement before and after the addition of ACh. The best-performing mutants were further confirmed and characterized after cloning into a mammalian expression vector, guiding further engineering efforts.

### Cell culture and transfection

HEK293A cells (Invitrogen) were cultured at 37 ℃ with 5% CO2 in Dulbecco’s Modified Eagle’s Medium (DMEM) with 10% fetal bovine serum (FBS), 2 mM L-glutamate, 100 U/mL penicillin and 100 μg/mL streptomycin. Cells were seeded at a density of 4.5 × 10^4^/cm^2^ in white 96-well tissue-culture plates (Greiner Bio-One) and transfected 24 h later with 50 ng plasmids or 1 μL of PCR reaction using Lipofectamine 3000 (Life Technologies) following the manufacturer’s instructions. Cells were assayed 24–48 h post-transfection.

### Primary neuronal culture and transfection

Cortical neurons were isolated from embryonic day 18 Sprague Dawley rat embryos by dissociation in RPMI medium containing 5 units/mL papain (Worthington Biochemical) and 0.005% DNase I at 37 °C and 5% CO2 in air. Dissociated neurons were plated at a density of 5.3 × 10^4^ cells/cm^2^ in white TC-treated 96-well plates with clear bottom (Corning) pre-coated overnight with > 300-kDa poly-D-lysine hydrobromide (Sigma-Aldrich). Cells were plated overnight in Neurobasal media with 10% FBS, 2 mM GlutaMAX, and B27 supplement (Life Technologies), then media were replaced with Neurobasal with 1% FBS, 2 mM GlutaMAX, and B27 the next day. Half of the media was replaced every 3–4 days with fresh media without FBS. 5-Fluoro-20-deoxyuridine (Sigma-Aldrich) was typically added at a final concentration of 16 mM at 7–9 DIV to limit glia growth. Cortical neurons were transfected at 9–11 DIV using a modified Lipofectamine 2000 (Life Technologies) transfection protocol in which media in each well of a 96-well plate was replaced for 60 min with 40 μL of DNA-lipid complexes (50 ng of anion exchange-purified ACh-NeuBI plasmids, and 0.1 μL of Lipofectamine 2000, in 40 μL of Neurobasal with 2 mM GlutaMAX per well). Neurons were assayed at 48 h post-transfection.

### Cell-based bioluminescence assay of indicators

ACh-NeuBI-expressing cells were assayed in Opti-MEM (for HEK293A cells) or Neurobasal media with 2 mM GlutaMAX (for cultured neurons) using Nano-Glo live cell assay system (Promega) per manufacturer’s instructions. Luminescence was measured on a Varioskan LUX multimode microplate reader (Thermo Scientific). Following baseline stabilization and signal plateau, the assay was temporarily paused for the addition of ACh or other neurotransmitters at predetermined concentrations. Subsequently, the plate was returned to the microplate reader to resume signal monitoring. Data analysis was done using signals from the last measurement prior to neurotransmitter addition with the peak signals recorded post-addition.

### In vitro characterization of purified indicators

pET-28a(+)-ACh-NeuBI0.1 plasmids were transformed into *E. coli* BL21(DE3) cells. Bacterial cultures were grown at 37 °C to reach an OD600nm of 0.6–0.8. Protein expression was induced using isopropyl β-D-1-thiogalactopyranoside (IPTG) at 16 °C for 16 h. Cells were harvested by centrifugation and lysed using TieChui™ *E. coli* Lysis Buffer (ACE Biotech) on ice. The recombinant proteins were purified using Ni-NTA affinity chromatography and were eluted using PBS (Sangon Biotech, E607008) containing 300 mM imidazole. Finally, the eluted proteins were desalted using a HiTrap desalting column (GE Healthcare) equilibrated with PBS (pH 7.4). For quantification of the purified protein indicators, absorbance at 280 nm was used to determine the protein concentration using the extinction coefficient calculated by the ExPASy’s ProtParam tool. Purified indicators were diluted in PBS and added to white 96-well plates. ACh was serially diluted and added to the wells to reach a final concentration as indicated. After incubation at 37 °C, luminescence was determined using the Nano-Glo Luciferase Assay System (Promega) following the manufacturer’s instructions with Varioskan ALF microplate reader (Thermo Scientific).

### Hydrodynamic transfection and bioluminescence imaging in mice

All animal procedures complied with USDA and NIH ethical regulations and were approved by the Stanford Institutional Animal Care and Use Committee according to protocols approved by the Administrative Panel on Laboratory Animal Care (APLAC) of Stanford University. Hydrodynamic transfection was conducted in 6- to 8-week-old male nude mice (NU/J, strain #: 002019, the Jackson Laboratory) by injecting 20 mg of pAAV-CMV-ACh-NeuBI or pAAV-CMV-ACh-NeuBImut plasmids in 2 mL saline via the tail vein within 5–6 seconds. 18–24 h after hydrodynamic transfection, bioluminescence imaging was performed by intraperitoneal (i.p.) injection of Dulbecco Phosphate Buffered Saline (DPBS, without Ca^2+^ or Mg^2+^, Corning) containing 0.93 mmol of FFz in each mouse. Images were acquired in a Lago-X optical imaging system (Spectral Instruments) every 30 seconds for 3–4 minutes, under 1.5–2.5% isoflurane in oxygen for anesthesia. Once the signals plateaued, the imaging was paused, and mice were i.p. injected with either ACh (10 mg/kg) or saline and put back into Lago-X optical imaging system to resume imaging. Imaging parameters were: open emission filter, 25-cm field of view, f/1.2 aperture, 2×2 binning, and 1–5 s exposure time depending on the brightness of the signals. Images were analyzed in Aura 4.0 software (Spectral Instruments). Data analysis was done using peak signals before and after the injection of ACh (10 mg/kg) or saline.

### Statistics

Student’s t-test and curve fitting by the “nonlinear regression (log(inhibitor) vs. response – variable slope)” model were performed in Prism 9 (GraphPad).

## Data availability

The main data supporting the findings of this study are available within the article and supplementary information. Additional raw data are available from the corresponding author upon request.

## Acknowledgements

We thank the Stanford Center for Innovation in In Vivo Imaging (SCi3) for use of instruments and assistance with in vivo bioluminescent imaging. We thank Yukun Alex Hao at Stanford for providing suggestions for the MIDAS screening. We thank Dr. Zi Yao at UCSF for reading the manuscript and offering valuable feedback. Illustrations were created with Biorender. Funding was provided by NIH grants 1R21NS122055 (M.Z.L.) and 1R21DA048252 (M.Z.L.), a Stanford Bio-X Interdisciplinary Initiatives Program Seed Grant (M.Z.L. and M.M.), and a Stanford Bio-X PhD Fellowship (Y.W.).

## Author contributions

Y.W. and M.Z.L. conceived of the project. Y.S. performed initial topology screening. Y.W. developed the MIDAS workflow and performed sensor optimization and characterization. Y.W., P.L., and L.X.L. performed animal experiments. Q.Q. characterized purified proteins. C.G. and M.H. synthesized and formulated luciferase substrates for animal imaging. M.Z.L, Y.S., and T.A.K. supervised the study. Y.W. and M.Z.L. wrote the manuscript with input from all authors.

